# Clinker: visualising fusion genes detected in RNA-seq data

**DOI:** 10.1101/218586

**Authors:** Breon M Schmidt, Nadia M Davidson, Anthony DK Hawkins, Ray Bartolo, Ian J Majewski, Paul G Ekert, Alicia Oshlack

## Abstract

Genomic profiling efforts have revealed a rich diversity of oncogenic fusion genes, and many are emerging as important therapeutic targets. While there are many ways to identify fusion genes from RNA-seq data, visualising these transcripts and their supporting reads remains challenging. Clinker is a bioinformatics tool written in Python, R and Bpipe, that leverages the superTranscript method to visualise fusion genes. We demonstrate the use of Clinker to obtain interpretable visualisations of the RNA-seq data that lead to fusion calls. In addition, we use Clinker to explore multiple fusion transcripts with novel breakpoints within the P2RY8-CRLF2 fusion gene in B-cell Acute Lymphoblastic Leukaemia (B-ALL).

**Availability and Implementation:** Clinker is freely available from Github https://github.com/Oshlack/Clinker under a MIT License.

## Introduction

Genomic structural abnormalities, such as translocations between and within chromosomes, are common in cancer and can result in the fusion of two genes which then function as an oncogenic driver. The first example of this was the recurrent t(9;22) fusion in Chronic Myeloid Leukaemia, creating the *BCR-ABL1* oncogene. This fusion gene results in a constitutively activated tyrosine kinase protein that can be effectively treated with small molecule inhibitors of ABL1, such as imatinib and dasatinib (Quintás-Cardama et al., 2009). The application of next generation sequencing in cancer, primarily transcriptome sequencing (RNA-seq), has subsequently identified thousands of different fusion genes in many cancer types (Mertens et al., 2015).

While there are many methods available for identifying fusion genes from RNA-seq data, there are few ways to visualise the fusion transcripts and the sequencing reads that support them. Simply aligning RNA-seq data to a reference genome or transcriptome does not allow clear visualisation of the translocation, or an appreciation of additional features such as splice variants. One visualisation strategy involves using predicted breakpoints to create the fusion transcript sequence, which can be used as a reference for read alignment (Beccuti et al., 2014). This approach demonstrates coverage across the fusion breakpoints but other information about the structure and expression of the fusion transcripts, such as its expression relative to non-fused transcripts, are lost. Some alternatives involve the construction of a new reference by joining the genomic sequence of the two genes involved in the fusion, or using the split screen view within IGV (Figure 2A) (Lågstad, 2017) (Haas and Tickle, 2017). However, because RNA-Seq read coverage is sparse in the genome, visualisation is hampered by the presence of introns.

**Figure 1.**
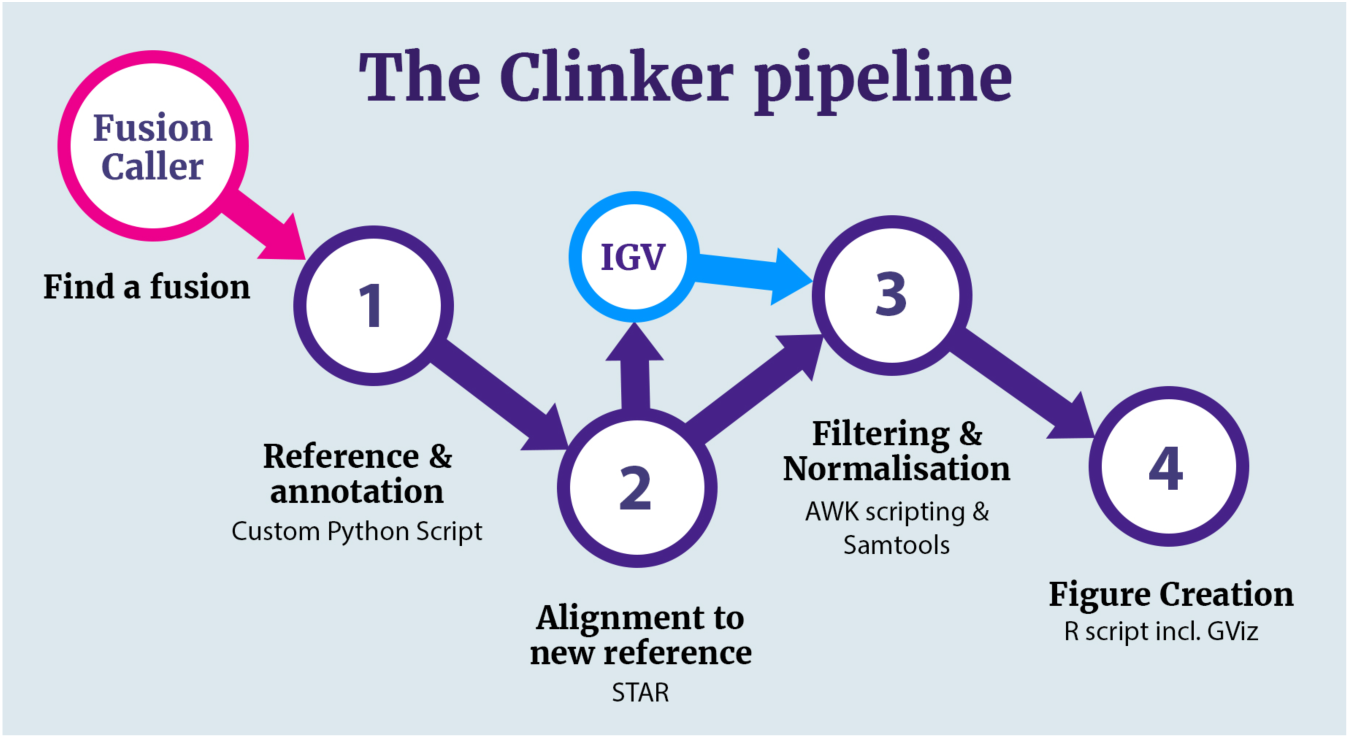
A visual representation of the Clinker pipeline. Users can choose to stop at step two, inspect fusion genes of interest in IGV, and then commence figure production for a refined list of fusion genes. The Fusion Caller (pink) and IGV (light blue) steps are external to Clinker.

**Figure 2.**
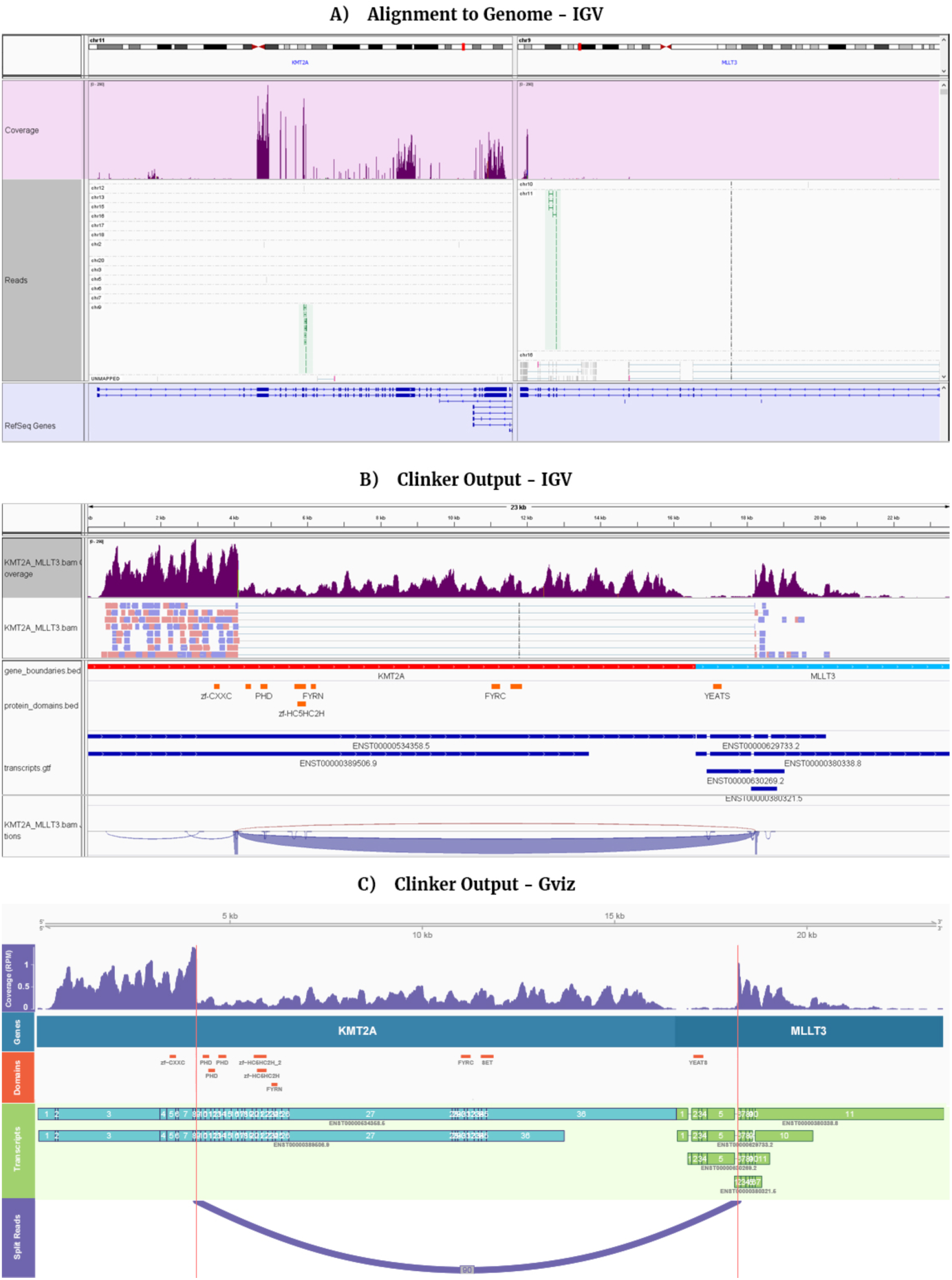
KMT2A-MLLT3 fusion gene visualised in IGV after alignment to the human genome (A). The backgrounds of the IGV tracks are coloured to distinguish between the coverage (purple), aligned reads (white) and annotation (blue), with green reads indicating that its partner is on a different chromosome. Such alignments may support the existence of a fusion. (B) Clinker output of the *KMT2A-MLLT3* gene fusion, visualised in IGV and (C) the GViz visualisation. The tracks in the Clinker GViz visualisation are (top to bottom): a superTranscript scale axis, a read coverage track, a gene boundary track, a protein domain track, a transcript (with exons annotation) track and a sashimi plot that indicates the fusion breakpoints (dark purple). The breakpoints are also indicated by the vertical lines. In addition to the Clinker tracks, the IGV visualisation includes a read support track.

With these considerations in mind we present Clinker, a tool for the visualisation, exploration and plotting of fusion genes found in RNA-seq data. Clinker utilises superTranscripts, a new type of transcriptome reference we developed that contains only the transcribed sequence of a gene, without introns, providing a highly compact reference for analysis and visualisation of RNA-Seq (Davidson *et al*., 2017). Clinker uses the human superTranscript references and creates fusion-superTranscripts by combining the two genes detected to be involved in a fusion event.

We have applied Clinker to a set of six B-cell ALLs that all report the *P2RY8-CRLF2* fusion to demonstrate several fusion isoforms. Clinker is a tool that provides direct visualization of fusion genes and an appreciation of their complexity that is not available using other methods.

## Materials & Methods

### Reference & Annotation Generation

The Clinker pipeline takes output from any fusion calling software, providing that it includes the hg19 or hg38 genomic coordinates of fusion gene breakpoints. These breakpoints are used to identify the two genes involved in the fusion and assigns them consistent gene symbols. This method is preferred even when gene symbols are provided by the fusion caller, due to the large variability in gene naming conventions. Once the two genes are identified, their sequences are retrieved from Clinker’s human superTranscript reference and concatenated to form a single fusion-superTranscript. An important feature of the fusion-superTranscript reference is that it includes the full sequence of both genes orientated in transcriptional direction. Thus, reads aligned to regions of the genes not involved in the fusion are also visualised, providing additional information about expression of these regions and the domains they encode. This is repeated for all fusion genes that have been identified in the sample. This results in a sample specific Clinker reference containing the fusion-superTranscripts, as well as the superTranscripts from all normal genes. We found it was important to map competitively to the non-fused genes in the reference to avoid spurious read alignments. Transcript, protein domain and gene boundary annotation files are also created using the Gencode24 hg38 reference (Harrow *et al*., 2012) and the Pfam protein database (Finn *et al*., 2016) to provide additional information for the visualisation.

### Alignment to the new reference

Clinker maps the sequencing reads to the newly generated reference using the STAR aligner (Dobin *et al*., 2013). The aligner must be splice aware as reads spanning the fusion breakpoint are identified as splice sites. The alignment stage of Clinker often yields greater read support for fusion genes than fusion callers. For example, in one sample JAFFA detected the *P2RY8-CRLF2* fusion with support from 53 spanning reads whereas Clinker, through STAR, reported 290 spanning reads (Table S1). As Clinker is given prior knowledge that a fusion exists between two genes, the fusion-superTranscript can be mapped to with less stringency, leading to the increase in successfully mapped reads across the breakpoint. After the alignment step, the mapped reads and fusion genes can be viewed with a genome viewer, such as IGV, by loading the Clinker reference FASTA, mapped reads, and the customised transcript, protein domain and gene boundary annotation tracks. IGV natively displays the fusion breakpoints and splice junctions through the splice junction track or sashimi plot (Figure 2B).

### Filtering, normalisation and figure creation

Once reads are aligned, filtering and normalisation steps are undertaken. Split reads with a small number of flanking bases on one side can be produced by incorrect split-read alignment. To account for this split reads with less than 5 base pairs of flanking sequence are immediately filtered out using both Samtools (Li *et al*., 2009) and custom AWK scripting (see Supplementary Figure 2 for an example). Coverage is normalised to reads per million (RPM) using STAR’s inbuilt normalisation function to allow comparison between samples.

A series of figures, one for each of the identified fusion genes, are then created using the R package, GViz (Hahne and Ivanek, 2016). These figures contain multiple tracks including coverage, gene boundaries, protein domains and the transcripts/exons involved in the fusion gene. A sashimi plot is also included in the figure to indicate the number of split reads that support the fusion, with three reads being set as a minimum threshold to further filter out spurious splicing events. The order, colour or presence of the tracks can be customised via the command line parameters of Clinker.

### Software requirements

Clinker can be run both manually and through Bpipe (Sadedin *et al*., 2012), a tool for running bioinformatics pipelines. The core dependencies for Clinker are STAR (Dobin *et al*., 2013), Samtools (Li *et al*., 2009) and Gviz (Hahne and Ivanek, 2016). Runtime was approximately 1 hour with 8 processors and 40 GB of memory allocated for a single publication quality figure and a further 1 minute for each additional figure. This test was conducted on a sample with approximately 130 million reads and 2007 fusion genes reported by JAFFA.

### P2RY8-CRLF2 cloning and expression

We have applied Clinker to a set of six B-cell ALL patient samples that all reported several non-canonical fusion isoforms within the *P2RY8-CRLF2* fusion gene. Fusion detection by RNAseq was confirmed by PCR using gene specific primers for *P2YR8* (5’-CAAGGTTGCTGGACAGATGGAA-3’) and *CRLF2* (5’-AATAGAGAATGTCGTCTCGCTGC-3’). Primers were designed to amplify products spanning the exons at the breakpoints of *P2RY8* and *CRLF2* in the mRNA transcripts detected by JAFFA. The alternate and frameshift fusions were cloned using primers to target the start of P2RY8 (5- CCCTGCACATGAGTGTTCAGAC-3’) and the end of *CRLF2* (5’- TCACAACGCCACGTAGGAG -3’), while the canonical fusion was amplified using a different *P2RY8* forward primer (5’- GCGGCCGCCTTTGCAAGGTTGC-3’). PCR products were cloned into P-GEM-T easy vector (Promega), Sanger sequenced and then subcloned into a retroviral pMSCV-GFP retroviral expression vector. Retrovirus was produced as previously described (Narayan *et al*., 2017), and transduced into IL3-dependant BaF3 cells. *CRLF2* was detected in the BaF3 cells using the Anti-Human TSLP receptor antibody (eBioscience) and the BD Cytofix/Cytoperm (BD Biosciences), according the manufacturer’s instructions. FACs analysis was performed on an LSRII flow cytometer (BD Biosciences).

## Results

### The Clinker pipeline

Clinker is an analysis pipeline that takes in fusion calls and raw RNA-seq reads and outputs a custom reference, mapped read data and image files to visualise and assess fusion transcripts. The steps in this pipeline are outlined in Figure 1 and described in detail in the Materials and Methods.

Briefly, before running Clinker, fusions are detected using one of the many specialised fusion gene callers, such as JAFFA (Davidson et al., 2015), STAR-fusion (Haas et al., 2017) or Pizzly (Melsted, 2017). Clinker proceeds by first concatenating the full-length superTranscripts of the two genes involved in the fusion and then adds them to a custom, sample specific superTranscriptome reference. Next, the reads are mapped back to the new reference using the STAR splice aware aligner (Dobin *et al*., 2013). A fusion can then be observed as splicing between the two concatenated genes. Finally, figures are generated that present the resulting splice junctions, coverage, protein domains and transcript annotation, for both the fusion and non-fused superTranscripts. Clinker outputs file formats that are compatible with IGV (Robinson *et al*., 2011), as well as publication quality images created with Gviz (Hahne and Ivanek, 2016).

### Clinker visualisations of reads, transcripts and protein domains

Most fusion calling algorithms use short-read RNA-seq data to report genes involved in potential fusion events as well as the number of reads detected that support these events. Figure 2 demonstrates the visualization of an *KMT2A-MLLT3* fusion gene that was detected in a B-cell ALL using JAFFA (Davidson *et al*., 2015) with and without using Clinker. Visualizing this fusion using IGV without Clinker is done using a split screen display of the regions of the genome spanning the fusion breakpoints (Figure 2A). While read pairs that span across the fusion breakpoints are viewable (green reads), the transcript context is difficult to discern. In contrast, using the Clinker superTranscript reference and outputs allows a neater and more informative visualization in IGV which can also display sashimi plots for the fusion support (Figure 2B). Finally Clinker also outputs a PDF image of the fusion that can be customised (Figure 2C).

### Identification of novel fusion isoforms in P2RY8-CRLF2

In order to demonstrate the utility of Clinker to provide visualization and insight into fusion genes we applied Clinker to six B-Cell Acute Lymphoblastic Leukaemia (B-ALL) samples that carried the *P2RY8-CRLF2* fusion. This fusion gene is reported to be present in ~7% of B-ALL cases and results in the overexpression of *CRLF2* (Mullighan *et al*., 2009). The canonical fusion joins the first non-coding (UTR) exon of P2RY8 to the coding region of *CRLF2* (Mullighan *et al*., 2009). Commonly, this fusion arises as a result of an interstitial deletion in the Par1 region of chrX or chrY (Mullighan *et al*., 2009). Interestingly, JAFFA called multiple breakpoints in *CRLF2* for this fusion gene in each of the sequenced B-ALL samples, suggesting different isoforms of this fusion. JAFFA identified the canonical break point in all samples. In addition, each of the six samples also expressed an isoform of *P2RY8-CRLF2* which joined the first exon of *P2RY8* to the 5’ UTR of *CRLF2*, and resulted in an in-frame transcript. This alternate fusion also featured the typical GT/AG donor/acceptor motif which exists at the majority of fusion junctions (Lai *et al*., 2015). The presence of the alternate breakpoints in the samples was confirmed using PCR and Sanger sequencing.

We used Clinker to visualize the *P2RY8-CRLF2* fusions in all six samples (Figure 3). Clinker detected the canonical breakpoint with the highest coverage in all samples (blue lines) and the novel 5’ breakpoint in all samples at much lower levels (red lines). Interestingly, all six samples had additional breakpoints between exon 1 and 2 of *CRLF2* (green lines) and one sample had a fourth breakpoint between exon 5 and 6 of *CRLF2* (yellow lines). However these additional transcripts are not in-frame.

**Figure 3.**
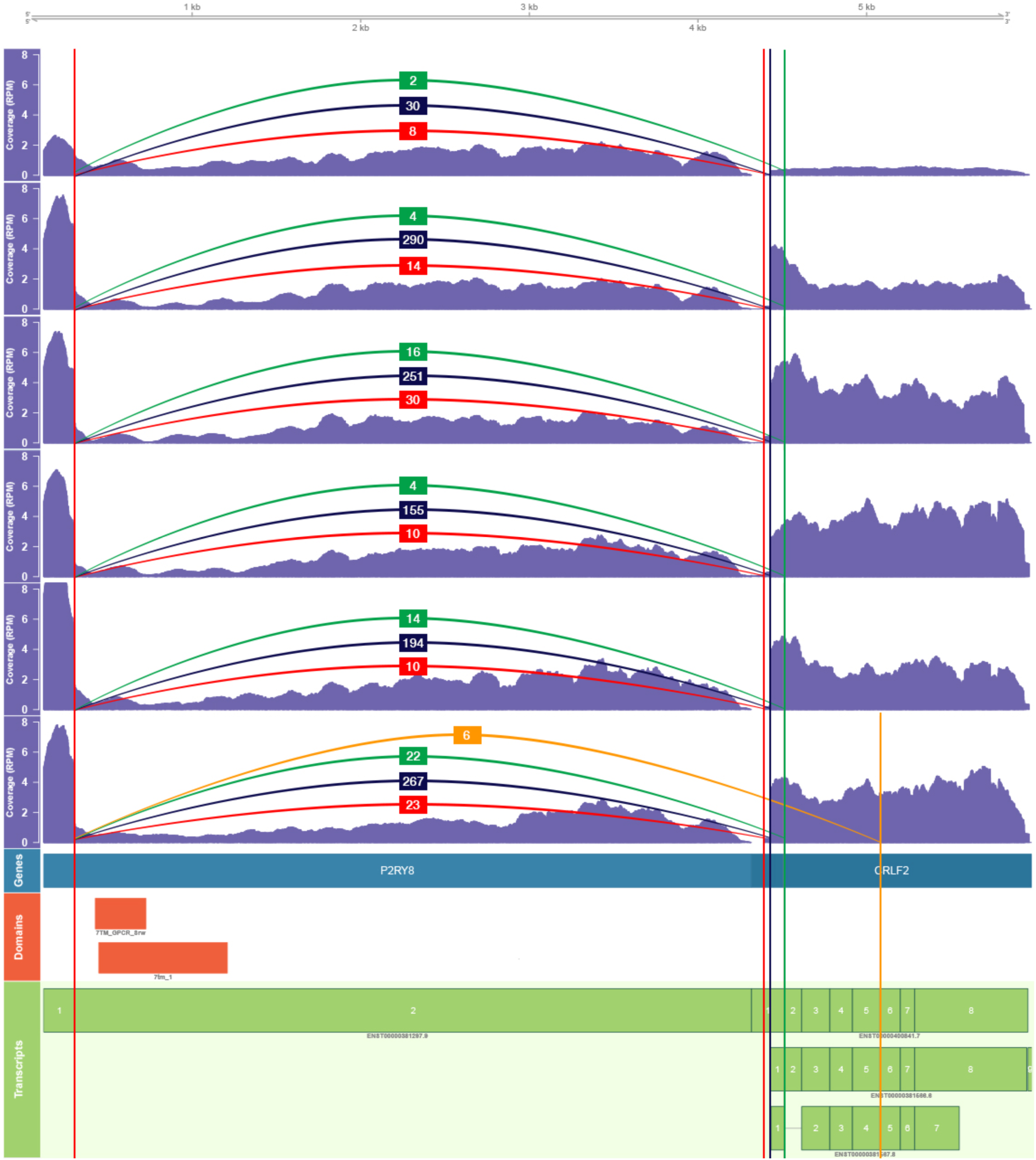
Visualisation of six samples containing the P2RY8-CRLF2 fusion. We combined the Clinker output (mapped reads, fusion superTranscript and annotation track) for the six samples using Gviz in R. From top to bottom: six coverage tracks with annotated breakpoints demonstrating read support, gene track, protein domains and gene transcripts. Each sample contains the canonical breakpoint (blue vertical line) as well as a novel upstream breakpoint occurring within the 5’UTR exon of CRLF2 (annotated with the red vertical line over the CRLF2 gene) along with two other breakpoints that are not in frame. The read support for this Clinker output can be compared to that of JAFFA’s in Supplementary Tables, 1 and 2.

To determine if the alternative, low abundance, in-frame transcript could drive *CRLF2* over expression, and so potentially contribute to the biology of ALL driven by *P2RY8-CRLF2* fusions we cloned the canonical and alternative version of the *P2RY8-CRLF2* fusion, as well as a frameshift version to act as a negative control, into retroviral vectors. The erythroleukaemia cell line (BaF3 cells) were then transduced with these retrorviruses to produce cell lines constitutively expressing the in-frame fusions or the negative (frameshift) *P2RY8-CRLF2* control. We measured *CRLF2* expression using an anti-human *CRLF2* antibody and flow cytometry (Supplementary Figure S3). The data show that both the canonical fusion and the alternate in-frame transcript can drive *CRLF2* overexpression in BaF3 cells, when compared to the frameshift control. These data suggest that the alternate transcript contributes to the overexpression of *CRLF2* in B-ALL.

## Conclusion

Here we present Clinker, a visualisation tool for exploring and plotting fusion genes discovered in RNA-seq data. Clinker allows raw reads involved in the discovery of fusion genes to be viewed and inspected in IGV and also generates annotations of transcripts and protein domains. This gives far greater insight into the expression levels and structures of the transcripts that make up the fusion gene. Furthermore, publication quality figures can be easily generated and refined using R functions. When we applied Clinker to real data we found that alternative breakpoints could be detected. Our examination of the *P2RY8-CRLF2* fusions indicates that these alternate forms exist in the primary samples, and may have biological relevance, as the alternate fusion appears capable of encoding a functional *CRLF2* protein.

